# Connexin 41.8 mediates the correct temporal induction of haematopoietic stem and progenitor cells

**DOI:** 10.1101/2023.07.27.550806

**Authors:** Tim Petzold, Masakatsu Watanabe, Julien Y. Bertrand

**Affiliations:** University of Geneva, Faculty of Medicine, Department of Pathology and Immunology, Rue Michel-Servet 1, Geneva 4, Switzerland; Graduate School of Frontier Biosciences, Osaka University, 1-3 Yamadaoka, Suita, Osaka 565-0871, Japan; Geneva Centre of Inflammation Research, University of Geneva, Faculty of Medicine, Rue Michel-Servet 1, Geneva 4, Switzerland

## Abstract

Haematopoietic stem and progenitor cells (HSPCs) derive from a subset of endothelial cells (ECs), known as haemogenic ECs by the process of endothelial-to-haematopoietic transition (EHT). Although many factors involved in EHT have been elucidated, we still have a poor understanding of the temporal regulation of this process. Mitochondrial-derived reactive oxygen species (ROS) have been shown to stabilise hypoxia-inducible factors 1/2α (Hif1/2α), allowing them to positively regulate EHT. Here, we show a developmental delay in EHT and HSPC induction in a gap junction mutant, *connexin (cx)41*.*8* (orthologous to mammalian *CX40*), in zebrafish. In mammalian cells, CX40 has been shown to localise to the mitochondria. We demonstrate that Cx41.8 is important for the correct temporal generation of mitochondrial ROS, which stabilise the Hif pathway, allowing for the subsequent specification of the haemogenic endothelium. Taken together, our data indicate that Cx41.8 mediates the correct induction of HSPCs.

## Introduction

HSPCs are rare, highly specialised cells which sit at the top of the haematopoietic hierarchy. HSPCs have the ability to self-renew and give rise to progenitor cells which differentiate into mature blood cells [1]. In vertebrates, HSPCs derive from the haemogenic endothelium [2], in a highly conserved process known as EHT [3-6]. Having a detailed understanding of all the factors involved in EHT may allow for *in vitro* generation of HSPCs from ECs in the future, which could have significant implications for regenerative medicine.

Connexin proteins play diverse roles in health and disease [7]. Six connexins form a connexon, which, when present at the cell membrane, can dock onto a connexon on a neighbouring cell to form a gap junction [8]. Gap junctions can facilitate the transport of ions, amino acids and small metabolites across the plasma membrane [9]. Interestingly, some connexins, such as CX40 in mammals, have been found to also localise to membranous intracellular organelles such as the mitochondria [10, 11]. CX40 has been shown to promote the production of ROS in mitochondria [10]. We previously demonstrated that zebrafish *cx41*.*8* (orthologous to mammalian *CX40*) played a role in HSPC expansion in the caudal haematopoietic tissue [12]. Indeed, HSPCs died by apoptosis in *cx41*.*8*^*t1/t1*^ mutants as a result of ROS toxicity during their expansion phase, whereas their specification and emergence were unaffected [12]. Here, we find that another *cx41*.*8* zebrafish mutant, *cx41*.*8*^*tq/tq*^, has a delay in the specification of the haemogenic endothelium resulting in delayed HSPC emergence. We determine that this phenotype is mechanistically linked with mitochondrial ROS production and the Hif pathway. We conclude that mitochondrial Cx41.8 contributes to the correct temporal induction of EHT and the subsequent formation of HSPCs during zebrafish development.

## Results

### *cx41*.*8*^*tq/tq*^ mutants harbour an HSPC specification defect

We previously characterized definitive haematopoiesis in the *cx41*.*8*^*t1/t1*^ mutant [12], and decided to investigate whether the *cx41*.*8*^*tq/tq*^mutant displays the same haematopoietic phenotype. The *leo*^*tq270*^mutant (referred to as *cx41*.*8*^*tq/tq*^throughout), possesses a missense mutation, I203F, in the fourth transmembrane domain of the protein [13] (Fig. 1 A-B), which results in disruption of the channel function [14]. Primitive haematopoiesis was found to be unaffected in *cx41*.*8*^*tq/tq*^embryos relative to controls, as determined by the expression of *gata1* and *pu*.*1* (primitive erythrocytes and primitive myeloid cells, respectively), by whole-mount *in situ* hybridisation (WISH) at 24hpf (Supp. Fig. 1 A-B). However, markers of definitive haematopoiesis were found to be significantly altered in *cx41*.*8*^*tq/tq*^ embryos: *gata2b* (an early marker of the haemogenic endothelium) expression at 24hpf (Fig. 1C), and *runx1* (a marker of nascent HSPCs) expression at 24hpf (Fig. 1D) and 28hpf (Supp. Fig.2A) were all reduced when compared with controls, as determined by WISH. In addition, *cmyb*^*+*^HSPCs budding from the dorsal aorta in *cmyb:GFP* embryos could not be observed in *cx41*.*8*^*tq/tq*^;*cmyb:GFP* embryos at 40hpf (Fig. 1E), providing further evidence for an HSPC specification defect in *cx41*.*8*^*tq/tq*^embryos.

**Figure 1.**
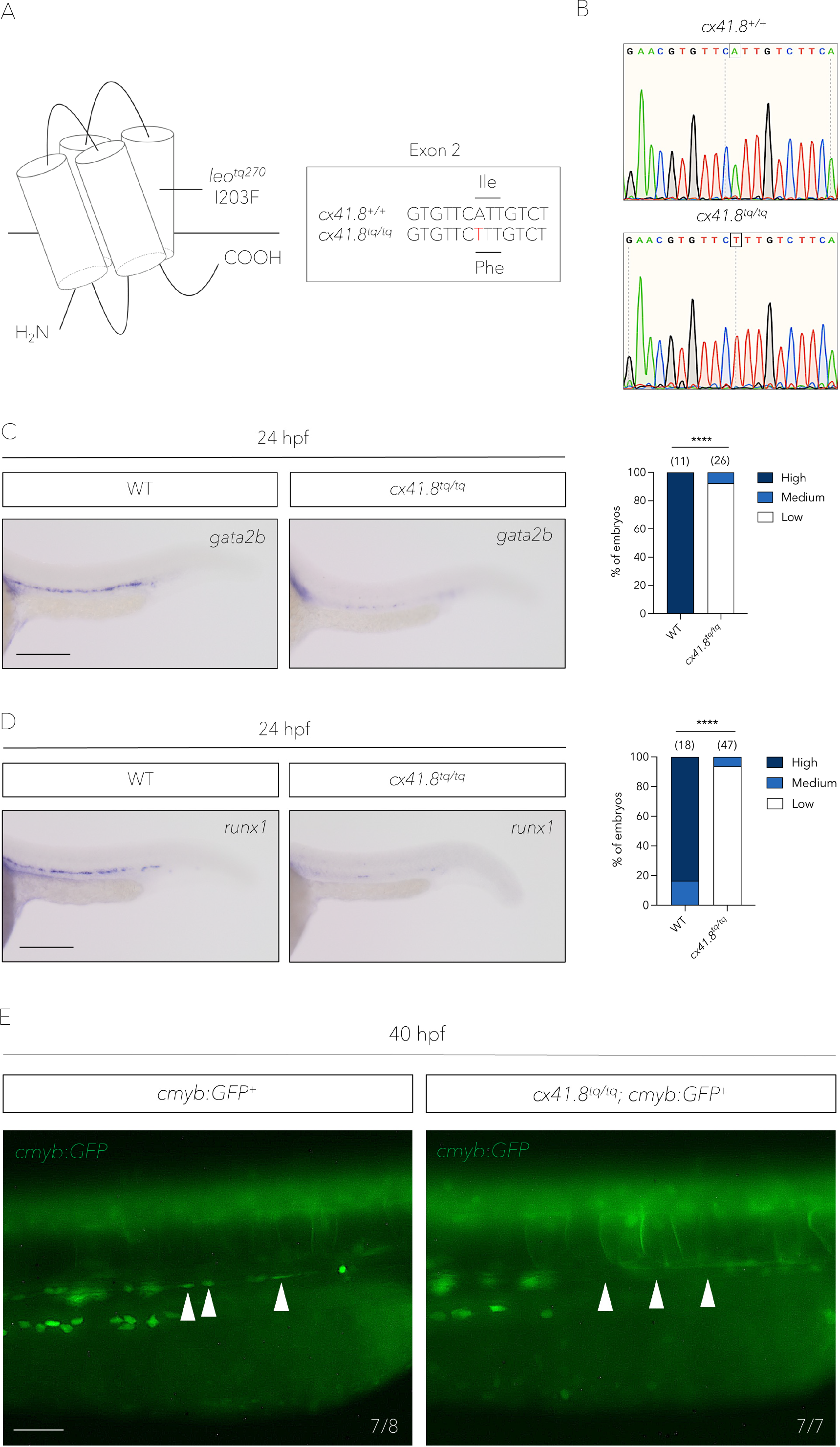
The I203F mutation in Cx41.8 results in a defect in haemogenic endothelium induction and HSPC specification. **A**. The *leo*^*tq270*^ (*cx41.8*^*tq/tq*^) mutant possesses an I203F change in the fourth transmembrane domain. **B**. Sanger sequencing shows an A-to-T base change in the *cx41.8*^*tq/tq*^ mutant. **C**. *in situ* hybridisation and quantification of *gata2b* in *cx41.8*^*tq/tq*^ mutants and controls at 24hpf. **D**. *in situ* hybridisation and quantification of *runx1* in *cx41.8*^*tq/tq*^ mutants and controls at 24hpf. **E**. Fluorescence imaging of dorsal aorta in 40hpf *cx41.8*^*tq/tq*^;*cmyb*:*GFP*^+^ embryos and controls (*cmyb*:*GFP*^+^). White arrowheads denote HSPCs budding from the dorsal aorta in *cmyb*:*GFP*^+^ embryos which are absent in *cx41.8*^*tq/tq*^;*cmyb*:*GFP*^+^ embryos. Numbers indicate the ratio of embryos with the respective phenotype. Statistical significance was calculated using a Chi-squared test. *p < 0.05, **p < 0.01, ***p < 0.001, ****p < 0.0001. Scale bars: 200 μm (**C** and **D**); 50 μm (**E**).

Since HSPCs are specified from the dorsal aorta in vertebrates [15], we asked whether arterial EC specification was impaired in *cx41*.*8*^*tq/tq*^embryos by analysing *dll4* expression. However, *dll4* expression was normal in *cx41*.*8*^*tq/tq*^embryos, as determined by WISH at 24hpf (Supp. Fig. 1C) and 28hpf (Supp. Fig. 1D). Altogether, this data indicates that *cx41*.*8* plays a role in the induction of the haemogenic endothelium and subsequent specification of HSPCs from the dorsal aorta.

### The HSPC specification defect in *cx41*.*8*^*tq/tq*^mutants is due to a delay in *gata2b* expression

To further characterise the HSPC specification defect present in *cx41*.*8*^*tq/tq*^mutants, we carried out WISH for the HSPC marker genes *runx1* and *cmyb* at later stages in development. HSPC numbers were found to be normal in *cx41*.*8*^*tq/tq*^ mutant embryos at 48hpf (Supp. Fig. 2B), 72hpf (Supp. Fig. 2C) and 4.5dpf (Supp Fig. 2D) as determined by *cmyb* WISH.

We therefore suspected a delay in the formation of the haemogenic endothelium in *cx41*.*8*^*tq/tq*^ mutants. To test this hypothesis, we determined the expression of *gata2b* at multiple stages in *cx41*.*8*^*tq/tq*^embryos, since *gata2b* expression precedes the expression of *runx1*, and controls the development of the haemogenic endothelium [16]. *gata2b* was found to be significantly reduced at 23hpf (Supp. Fig. 3A), 24hpf (Supp. Fig. 3B), 26hpf (Supp. Fig. 3C) and 28hpf (Supp. Fig. 3D) in *cx41*.*8*^*tq/tq*^ mutants, compared to control embryos. However, *cx41*.*8*^*tq/tq*^embryos displayed significantly more *gata2b* expression at 30hpf, 32hpf and 36hpf (Supp. Fig. 4A-C), whilst at 48hpf (Supp Fig. 4D), no difference in *gata2b* expression was present. This data demonstrates delayed *gata2b* expression in the dorsal aorta of *cx41*.*8*^*tq/tq*^embryos, which we speculate also results in the delay in the expression induction of the downstream genes, *runx1* and *cmyb*. As such, *cx41*.*8*^*tq/tq*^embryos possess a developmental delay in the formation of the haemogenic endothelium, which subsequently results in the disrupted temporal control of HSPC specification.

### *cx41.8* is expressed in ECs in the dorsal aorta floor

Our lab has previously demonstrated that *cx41.8* is expressed in vascular ECs in zebrafish [17], and we also showed a critical role for Cx41.8 in bridging HSPCs to their vascular niche in the caudal haematopoietic tissue [12]. Additionally, transcriptomics data recently revealed a high expression of *cx41.8* in arterial ECs at 24hpf [18]. In order to investigate the spatiotemporal expression pattern of *cx41.8* more precisely, we generated a *cx41.8:EGFP* zebrafish reporter line using the previously described *cx41.8* promoter [13] (Fig. 2A).

**Figure 2.**
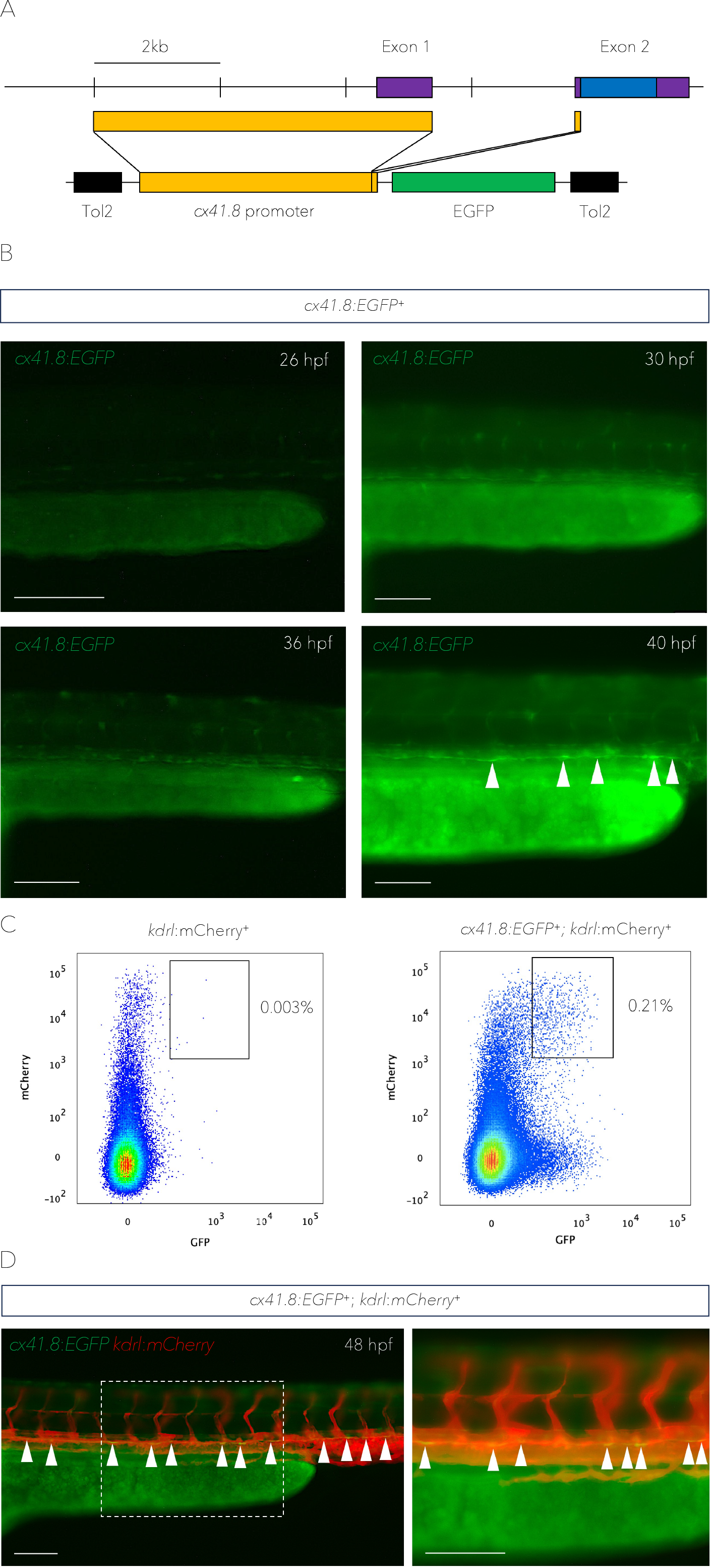
*cx41.8* is expressed in presumptive haemogenic endothelial cells of the dorsal aorta. **A**. Design of the *cx41.8*:*EGFP* plasmid. The upper line indicates the *cx41.8* locus structure. The lower line indicates the construct design. Purple boxes indicate the *cx41.8* exons; blue boxes indicate the open reading frame; yellow boxes indicate the 4.5kb sequence upstream of the *cx41.8* start codon; black boxes indicate the transposon sequences and the green box indicates the EGFP coding sequence. **B**. *cx41.8:EGFP* expression in presumptive vasculature including the aorta from 26-40hpf. White arrowheads denote presumptive haemogenic endothelial cells in the floor of the aorta. **C**. Flow cytometry analysis of double-positive cells in 48hpf *kdrl*:*mCherry*^+^ or *cx41.8*:*EGFP*^+^; *kdrl*:*mCherry*^+^ embryos. **D**. Expression of *cx41.8*:*EGFP* and *kdrl*:*mCherry* at 48hpf. White arrowheads denote *cx41.8*:*EGFP* and *kdrl*:*mCherry* double-positive endothelial cells in the floor of the dorsal aorta. Scale bars: 100 μm (**B** and **D**).

During HSPC specification (26-54hpf), *EGFP* was expressed in structures resembling vasculature. In particular, strong expression was detected as a thin line in the region of the axial vasculature (likely the aortic floor (Fig. 2B)). To confirm these findings, we generated *cx41.8:EGFP;kdrl:mCherry* double transgenic embryos. The region containing the trunks and tails were dissected at 48hpf and subjected to flow cytometry analyses. A population of double positive *cx41.8:EGFP;kdrl:mCherry* cells was detected, which is absent in *kdrl:mCherry* embryos (Fig. 2C, for gating strategy see Supp. Fig. 5). This confirms that *cx41.8* is expressed in ECs of the zebrafish trunk and tail.

Finally, by observing *cx41.8:EGFP;kdrl:mCherry* embryos by fluorescence microscopy, we confirmed that double positive *cx41.8:EGFP;kdrl:mCherry* ECs are indeed present in the floor of the aorta (Fig. 2D). Together, this demonstrates that *cx41.8* is expressed in presumptive haemogenic ECs in the aortic floor during the time of EHT and HSPC specification.

### Temporal mitochondrial ROS induction of the HSPC program requires *cx41.8*

Previous work has elucidated a key role of mitochondrial-derived ROS in HSPC specification [19]. Furthermore, the mammalian Cx41.8 orthologue, CX40, has been found to be localised to the mitochondria in mouse and human ECs, and was shown to be necessary for mitochondrial ROS production [10]. Hence, we wanted to test if the production of mitochondrial ROS in aortic floor haemogenic ECs was impaired in *cx41.8*^*tq/tq*^ mutants. To determine whether this was the case, we first probed for the presence of total cellular ROS and mitochondrial-derived ROS in the aorta at 16hpf (Fig. 3A). Total cellular ROS were detected in the ventral side of the vascular chord in *kdrl:GFP* embryos, but were absent in *cx41.8*^*tq/tq*^;*kdrl:GFP* embryos, as determined by using a cellROX probe (Fig. 3B). Similarly, mitochondrial ROS were also detected on the ventral side of the vascular chord in *kdrl:GFP* embryos at 16hpf, but were absent in *cx41.8*^*tq/tq*^;*kdrl:GFP* embryos, as determined using a mitoSOX probe (Fig. 3C). This indicates that there is indeed a defect in mitochondrial ROS production in aortic ECs in *cx41.8*^*tq/tq*^ mutant embryos, prior to the induction of *gata2b* expression.

**Figure 3.**
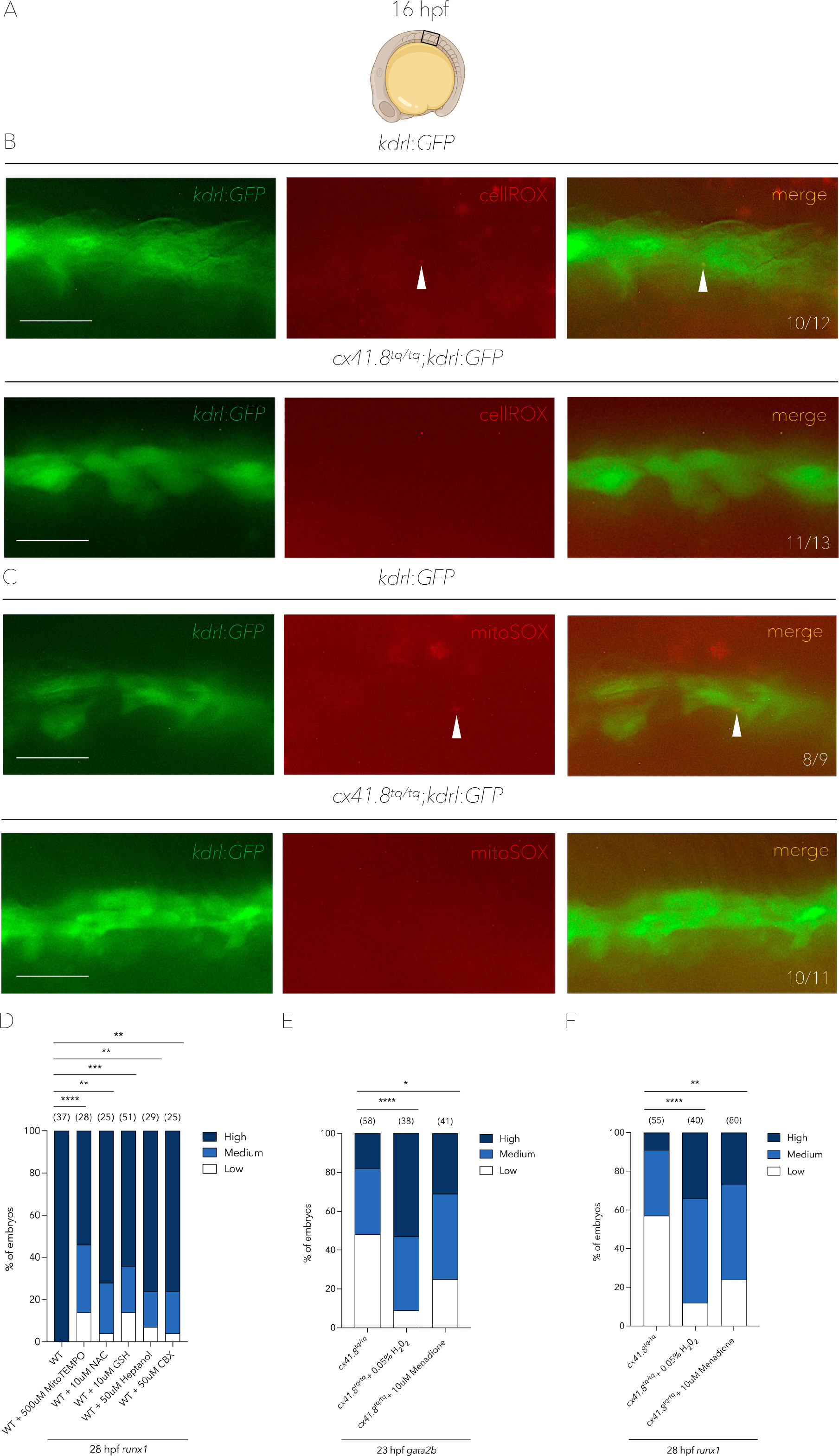
Mitochondrial-derived ROS production in endothelial cells is required for haemogenic endothelium induction and the specification of HSPCs. **A**. Schematic showing the region of 16hpf embryos which was analysed by fluorescence microscopy in **B** and **C. B**. Fluorescence microscopy images of total cellular ROS detection in *kdrl*:*GFP*^+^ or *cx41.8*^*tq/tq*^;*kdrl*:*GFP*^+^ embryos. White arrowheads denote the presence of ROS in endothelial cells. Numbers indicate the ratio of embryos with the respective phenotype. **C**. Fluorescence microscopy images of mitochondrial-derived ROS detection in *kdrl*:*GFP*^+^ or *cx41.8*^*tq/tq*^;*kdrl*:*GFP*^+^ embryos. White arrowheads denote mitochondrial-derived ROS in endothelial cells. Numbers indicate the ratio of embryos with the respective phenotype. **D**. Quantification of aortic *runx1* signal (*in situ* hybridisation) at 28hpf in WT control embryos and those treated with MitoTEMPO, NAC, GSH, heptanol or CBX. **E**. Quantification of aortic *gata2b* signal (*in situ* hybridisation) at 23hpf in *cx41.8*^*tq/tq*^ control embryos and *cx41.8*^*tq/tq*^ embryos treated with H_2_0_2_ or menadione. **F**. Quantification of aortic *runx1* signal (*in situ* hybridisation) at 28hpf in *cx41.8*^*tq/tq*^ control embryos and *cx41.8*^*tq/tq*^ embryos supplemented with H_2_0_2_ and menadione. Statistical significance was calculated using a Chi-squared test. *p < 0.05, **p < 0.01, ***p < 0.001, ****p < 0.0001. Scale bars: 25 μm (**B** and **C**).

Next, we set out to determine whether modulation of ROS has an effect on haemogenic endothelium induction and HSPC specification. Wild-type (WT) embryos were treated with MitoTEMPO, a specific inhibitor of mitochondrial ROS production [20], and subsequently fixed for *in situ* hybridisation. Treatment with MitoTEMPO from 14hpf resulted in impaired HSPC specification as determined by a significant reduction of *runx1* WISH signal at 28hpf (Fig. 3D). Treatment of WT embryos with the anti-oxidants N-Acetyl-Cysteine (NAC) or reduced L-glutathione (GSH) from 14hpf, also impaired HSPC specification as determined by *runx1* WISH at 28hpf (Fig. 3D), as was previously shown [19]. Moreover, treatment of WT embryos with the connexin blockers heptanol [21] and carbenoxolone (CBX) [22] from 14hpf also resulted in a decrease in HSPC specification (Fig. 3D).

Following this, we treated *cx41.8*^*tq/tq*^ mutant embryos with the ROS enhancers, H_2_O_2_ [23] and menadione [24] from 14hpf, which resulted in an increased expression of both *gata2b* at 23hpf (Fig. 3E) and *runx1* at 28hpf (Fig. 3F), as determined by WISH. H_2_O_2_ was able to rescue *gata2b* expression in a dose-dependent manner (Supp. Fig. 6). Together, this data indicates that Cx41.8 plays a key role for the correct temporal generation of mitochondrial ROS and the subsequent induction of the haemogenic program in the dorsal aorta.

### Induction of the Hif1/2α-mediated haematopoietic program in response to mitochondrial ROS is dependent on *cx41.8*

Recent research has demonstrated that hypoxia and mitochondrial ROS are required for the stabilisation of the transcription factors Hif1/2α [19, 25]. Mechanistically, ROS stabilises Hif1/2α by inhibiting prolyl hydroxylases which target Hif1/2α for ubiquitination by the von Hippel Lindau (VHL) protein, resulting in their subsequent degradation [26, 27]. Hif1/2α have been shown to act upstream of Notch1a/b signalling, which in turn induces *gata2b* expression and the formation of the haemogenic endothelium [25]. Therefore, we hypothesized that the lack of mitochondrial ROS production at 16hpf in *cx41.8*^*tq/tq*^ embryos resulted in the degradation of Hif1/2α. and the subsequent lack of *gata2b* transcriptional activation. To test whether *cx41.8* is involved in this ROS-Hif1/2α-Notch1a/b-*gata2b* pathway, *cx41.8*^*tq/tq*^ embryos were treated with either cobalt chloride (CoCl_2_), a hypoxia mimetic which interferes with the interaction between Hif1/2α and *vhl* [28], or the prolyl hydroxylase inhibitor dimethyloxallyl glycine (DMOG) [29]. Treatment with CoCl_2_ or DMOG from 14hpf resulted in a rescue of both *gata2b* (Fig. 4A) and *runx1* (Fig. 4B) expression in *cx41.8*^*tq/tq*^ embryos at 23 and 28hpf, respectively.

**Figure 4.**
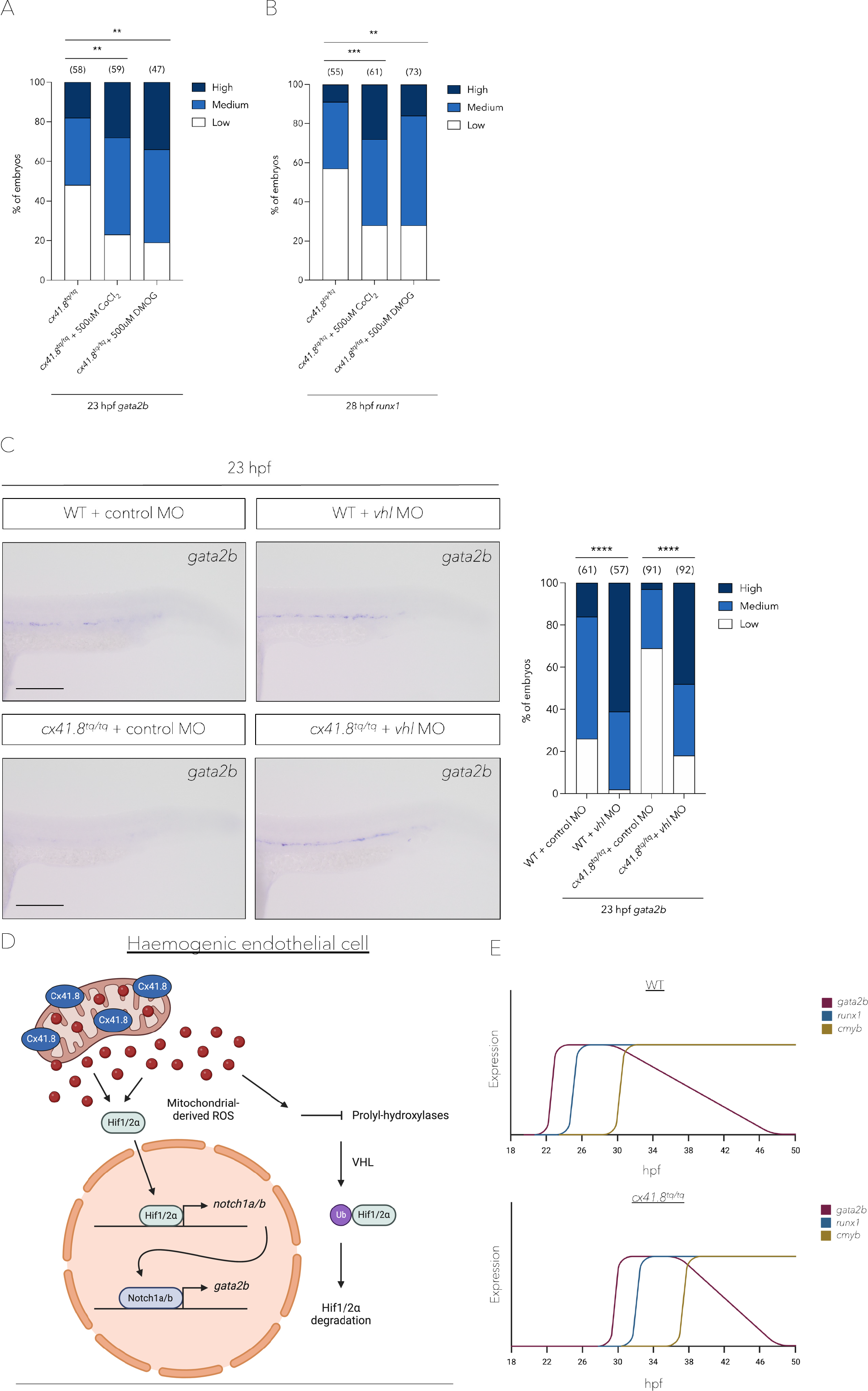
Stabilisation of Hif1/2α rescues haemogenic endothelium induction and HSPC specification in *cx41.8*^*tq/tq*^ mutants. **A**. Quantification of aortic *gata2b* signal (*in situ* hybridisation) at 23hpf in *cx41.8*^*tq/tq*^ control embryos and *cx41.8*^*tq/tq*^ embryos supplemented with CoCl_2_ or DMOG. **B**. Quantification of aortic *runx1* signal (*in situ* hybridisation) at 28hpf in *cx41.8*^*tq/tq*^ control embryos and *cx41.8*^*tq/tq*^ embryos treated with CoCl_2_ or DMOG. **C**. *in situ* hybridisation and quantification of *gata2b* at 23hpf in WT control embryos and *cx41.8*^*tq/tq*^ embryos injected with either control- or *vhl*-MO. **D**. Working model to represent how Cx41.8 localises to the mitochondria in haemogenic endothelial cells, allowing mitochondrial ROS production which stabilises Hif1/2α, which in turn induces *gata2b* expression via Notch1a/b signalling. **E**. Depiction of differences in the timing of expression of the HSPC program genes *gata2b, runx1* and *cmyb* in WT and *cx41.8*^*tq/tq*^ embryos. Statistical significance was calculated using a Chi-squared test. *p < 0.05, **p < 0.01, ***p < 0.001, ****p < 0.0001. Scale bars: 100 μm (**C**).

Finally, we used a previously described *vhl* morpholino (MO) [30-32] to prevent *vhl* function. The *vhl-*MO resulted in the induction of a cryptic splice site in exon 1 of the *vhl* transcript (Supp. Fig. 7A) and the subsequent loss of 18 amino acids from the VHL beta domain (Supp. Fig. 7B) which is required for the interaction between VHL and Hif1/2α [33]. MO-mediated knockdown of *vhl* in WT embryos resulted in an increase in *gata2b* expression at 23hpf (Fig 4C). Furthermore, MO-mediated knockdown of *vhl* in *cx41.8*^*tq/tq*^ embryos also rescued *gata2b* expression at 23hpf (Fig 4C) and *runx1* expression at 28hpf (Supp. Fig. 8), showing that the delay in *gata2b* expression was likely due to the degradation of Hif1/2α.

## Discussion

Here, we have demonstrated that there is a delay in the development of the haemogenic endothelium and the subsequent specification of HSPCs in the absence of a functional Cx41.8. Mechanistically, Cx41.8 seems to play a key role in the production of mitochondrial ROS. This may be the result of reduced calcium uptake into the mitochondria in *cx41.8*^*tq/tq*^ mutants, as has been reported in *CX40*-deficient ECs [10]. Furthermore, whilst an increase in mitochondrial metabolism was previously found to drive mitochondrial ROS generation in response to glucose, enhancing HSPC induction [19], calcium entry into the mitochondria has also been demonstrated to induce mitochondrial metabolism [34, 35]. Hence, we speculate that mitochondrial calcium influx, occurring in a Cx41.8 dependent manner, drives mitochondrial metabolism, resulting in an increase in mitochondrial ROS production. Ultimately, this elevation in ROS induces the downstream haematopoietic program.

The partial functionality of the Cx41.8 channel in *cx41.8*^*tq/tq*^ mutants [14] may explain why the HSPC program is eventually induced. However, this could also result from functional redundancy between Cx41.8 and other connexins such as Cx43 or Cx45.6 in the mitochondria, since they are also expressed in zebrafish arterial ECs at 24hpf [18] and *cx43* knockdown has previously been shown to result in an HSPC specification defect in zebrafish [36]. This potential functional redundancy may also provide an explanation as to why HSPCs are specified normally, without any delay, in *cx41.8*^*t1/t1*^ embryos [12]. In these null mutants, *cx41.8* expression is completely absent but may be functionally compensated by other connexins, whereas in *cx41.8*^*tq/tq*^ mutants, although *cx41.8* is expressed, its channel function is reduced [14]. Moreover, as Cx41.8 may form heterotypic channels with Cx43 and/or Cx45.6 (and potentially also with others), the function of these chimeric channels would also be altered.

In summary, we suggest that mitochondrial channels formed by Cx41.8 and perhaps also others, are important for ROS production in the mitochondria of dorsal aorta ECs, as early as 16hpf. These mitochondrial-derived ROS stabilise the transcription factors Hif1/2α, which subsequently translocate into the nucleus leading to the expression induction of *gata2b* in a Notch1a/b-dependent manner (Fig. 4D), as shown previously [25]. This signalling cascade ends with the specification of the haemogenic endothelium, leading to the birth of HSPCs. *cx41.8*^*tq/tq*^ mutants display a delay in *gata2b, runx1* and *cmyb* expression (Fig. 4E). Hence, *cx41.8* facilitates the correct temporal induction of HSPCs. We speculate that this is likely to also be the case in mammals, since *CX40* (the mammalian orthologue of *cx41.8*) is highly expressed in the haemogenic endothelium of mouse [37] and humans [38], and as CX40 localises to the mitochondria in ECs of both these species [10].

Further research will be required to determine the exact mechanism(s) by which mitochondrial ROS are produced specifically in this EC subpopulation. This may involve sterile inflammation, which is particularly important for the induction of EHT and the budding of nascent HSPCs from the aortic floor [39-41]. Our data contribute to a better understanding of the factors required for the initiation of the haemogenic program, which may have significant implications for enhancement of current regenerative medicine protocols, to produce haematopoietic progenitor cells *in vitro*.

## Methods

### Ethical statement

Zebrafish were raised in accordance to FELASA and Swiss guidelines [42]. No authorisation was required since experiments were carried out on embryos up to 5 days post fertilization. All efforts were made to comply to the 3R guidelines.

### Zebrafish husbandry

AB* zebrafish, as well as transgenic zebrafish lines were kept in a 14/10 hr light/dark cycle at 28.5°C. Embryos were obtained as described previously [43]. Embryos were staged by hours-post fertilization (hpf) as described previously [44]. In this study the mutant zebrafish line *leo*^*270/270*^[14] (referred to as *cx41.8*^*tq/tq*^) was utilised and genotyped by PCR (using primers listed in Supp. Table. 1), followed by Sanger sequencing (Fig. 1B). The following transgenic lines were used in this study: *Tg(kdrl:GFP)*^*s843*^[45], *Tg(kdrl:Has.HRASmCherry)*^*s896*^ [46] (referred to as *Tg(kdrl:mCherry)*), *Tg(cmyb:GFP)*^*zf169*^[47] and *Tg(cx41.8:EGFP)* [13]. Zebrafish embryos were treated with 0.003% 1-phenyl-2-thiourea (PTU, Sigma P7629) starting at 24hpf to prevent pigmentation.

### Generation of transgenic animals

For *Tg(cx41.8:EGFP)* zebrafish generation, 50pg of the Tol2 *cx41.8:EGFP* plasmid, described previously [13], was co-injected with 50pg of *tol2 transposase* mRNA into AB* zebrafish embryos. Injected F0s were mated with AB* zebrafish, and the resulting F1 offspring were screened by fluorescence microscopy to assess germline integration of the Tol2 construct. F2 zebrafish adults were subsequently mated and their offspring utilised in experiments.

### Whole-mount *in situ* hybridisation

Whole-mount *in situ* hybridisation (WISH) was performed on 4% paraformaldehyde-fixed embryos as described previously [48]. Digoxigenin-labelled *dll4, gata1, pu.1, gata2b, runx1* and *cmyb in situ* probes were used and their generation has been described previously [12, 49].

### Chemical treatments

All compounds used in this study were purchased from Sigma-Aldrich. Zebrafish embryos were exposed to compounds in 0.003% 1-phenyl-2-thiourea (PTU, Sigma, P7629) E3 (fish) water in multi-well plates from 14hpf to either 23 or 28hpf. Following exposure, embryos were fixed in 4% paraformaldehyde. All chemical treatment experiments are a combination of at least two independent experiments with independent clutches.

### Flow cytometry

Dissected embryos were incubated with a liberase-blendzyme 3 (Roche) solution for 90min at 33 °C, then dissociated and resuspended in 0.9x PBS-1% fetal calf serum, as described previously [50]. We distinguished and excluded dead cells by staining them with SYTOX Red (Life Technologies). Cell suspensions were passed through a 40mm filter prior to flow cytometry. Data were acquired on a LSR2Fortessa (BD Biosciences, software diva8.0.2) and analysed with FlowJo (v10).

### Total cellular and mitochondrial reactive oxygen species detection

Whole-mount staining with the cellROX (Invitrogen) or MitoSOX (Life Technologies) probes was performed on living zebrafish embryos at 16hpf, following the methods described previously [51]. Embryos were exposed to either a 5 μM cellROX or a 5 μM MitoSOX solution for 45 minutes and incubated at 28.5°C. Subsequently, imaging was carried out by fluorescence microscopy.

### Microscopy

Whole mount *in situ* hybridisation images were taken on an Olympus MVX10 microscope in 100% glycerol. Fluorescent images were taken with an IX83 microscope (Olympus). All images were taken using the CellSens Dimension software (Olympus).

### Morpholino injections

The standard control (CCTCTTACCTCAGTTACAATTTATA) and *vhl* (GCATAATTTCACGAACCCACAAAAG) morpholino (MO) oligonucleotides were purchased from GeneTools (Philomath, OR). MO efficiency was tested by PCR (Supp. Fig. 7C) from total RNA extracted from 15 embryos per sample at 24hpf (using primers listed in Supp. Table. 2). The *vhl*-MO-induced loss of 54 bp from exon 1 of the *vhl* transcript upon injection of 6 ng of MO was confirmed by Sanger sequencing (Supp. Fig. 7D). In all subsequent MO experiments, 6 ng of *vhl* or standard control MO was injected per embryo. All morpholino experiments are a combination of 3 experiments with independent clutches.

### Image processing and WISH phenotypic analyses

All images were processed using Fiji ImageJ (NIH) [52]. WISH phenotypic variation was analysed qualitatively and depicted graphically as the percentage of total embryos scored exhibiting high, medium or low gene expression in the region of interest before statistical analyses was carried out.

## Data analyses

At least three independent experiments were carried out in all cases, unless stated otherwise. In all experiments, normality was assumed and variance was comparable between groups. Sample size was selected empirically according to previous experience in the assessment of experimental variability. Numerical data are the mean ± s.e.m., unless stated otherwise. For WISH analyses, statistical significances were calculated using either a Chi-squared or Fisher’s test, where appropriate (denoted as *p < 0.05, **p < 0.01, ***p < 0.001, ****p < 0.0001 in all figures). Statistical calculations and the graphs for the numerical data were performed using Prism 9 software (GraphPad Software).

## Data Availability

All raw data will become freely accessible on Yareta following publication.

## Author contribution

T.P. performed all experiments and analyses. T.P. and J.Y.B. designed experiments. M.W. generated the Tol2 plasmid containing the *cx41.8:EGFP* reporter. T.P. generated the *cx41.8:EGFP* reporter line. T.P. and J.Y.B. wrote the original draft of the manuscript. T.P., M.W. and J.Y.B. edited the manuscript.

## Acknowledgements

We would like to thank all lab members for their comments and suggestions during this project. We would also like to thank Prof. Brenda Kwak for helpful discussions about connexins. T.P. benefitted from a grant from the Gabbiani Fund. J.Y.B. was funded by the Swiss National Fund (grant #310030_184814) and by the Fondation Privée des Hopitaux de Genève. Fig. 1A, Fig. 4D and zebrafish figures were all created using biorender.com.

