## supplemental material for "Connexin 41.8 mediates the correct temporal induction of haematopoietic stem and progenitor cells"

#### Supplementary Information

| Mutant | Forward | Reverse |
| --- | --- | --- |
| *cx41.8*$^{tq/tq}$ | TGCTGCAAACATACGTCCTC | TTTGCAGAGTTCTGCTGGTG |

**Supplementary Table 1. Primers used for genotyping *cx41.8*$^{tq/tq}$ mutant embryos**

| Gene | Forward | Reverse |
|---|---|---|
| *vhl* | CAGGTCAACGTTCTGTTCTG | GTGATCTTGGCATTTGCGAC |

**Supplementary Table 2. Primers used to analyse the *vhl* knockdown efficiency of the *vhl*-MO**

Supplementary Figure 1

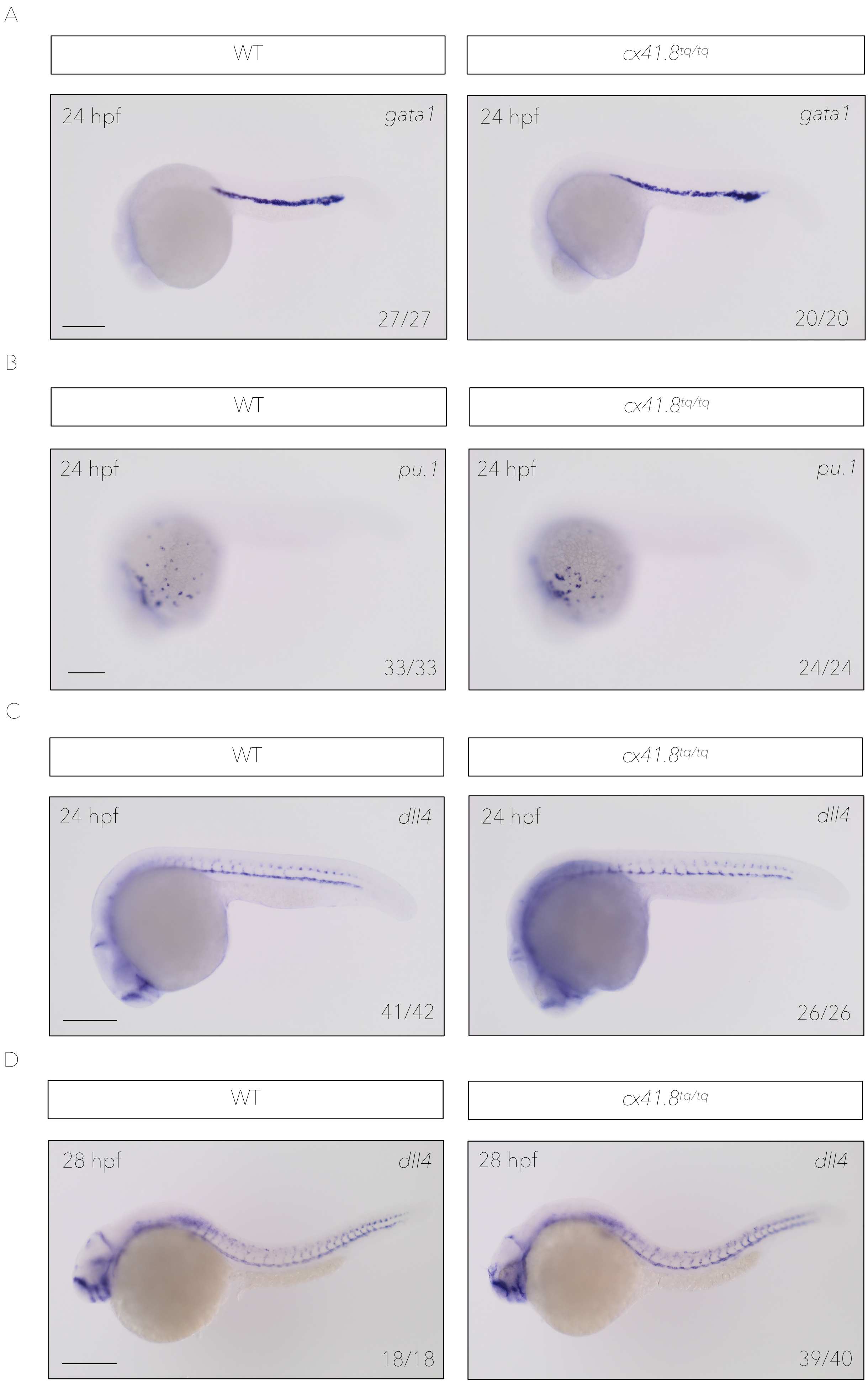

**Supplementary Figure 1. *cx41.8$^{tq/tq}$* mutant embryos do not have altered primitive haematopoiesis or vascular development**

**A.** *in situ* hybridisation against *gata1* (primitive erythrocytes) in *cx41.8$^{tq/tq}$* mutants and controls at 24hpf. **B.** *in situ* hybridisation against *pu.1* (primitive macrophages) in *cx41.8$^{tq/tq}$* mutants and controls at 24hpf. **C.** *in situ* hybridisation against *dll4* (arterial endothelium) in *cx41.8$^{tq/tq}$* mutants and controls at 24hpf. **D.** *in situ* hybridisation against *dll4* in *cx41.8$^{tq/tq}$* mutants and controls at 28hpf. Numbers indicate the ratio of embryos with the respective phenotype. Scale bars: 200 μm (**A**, **B**, **C** and **D**).

Supplementary Figure 2

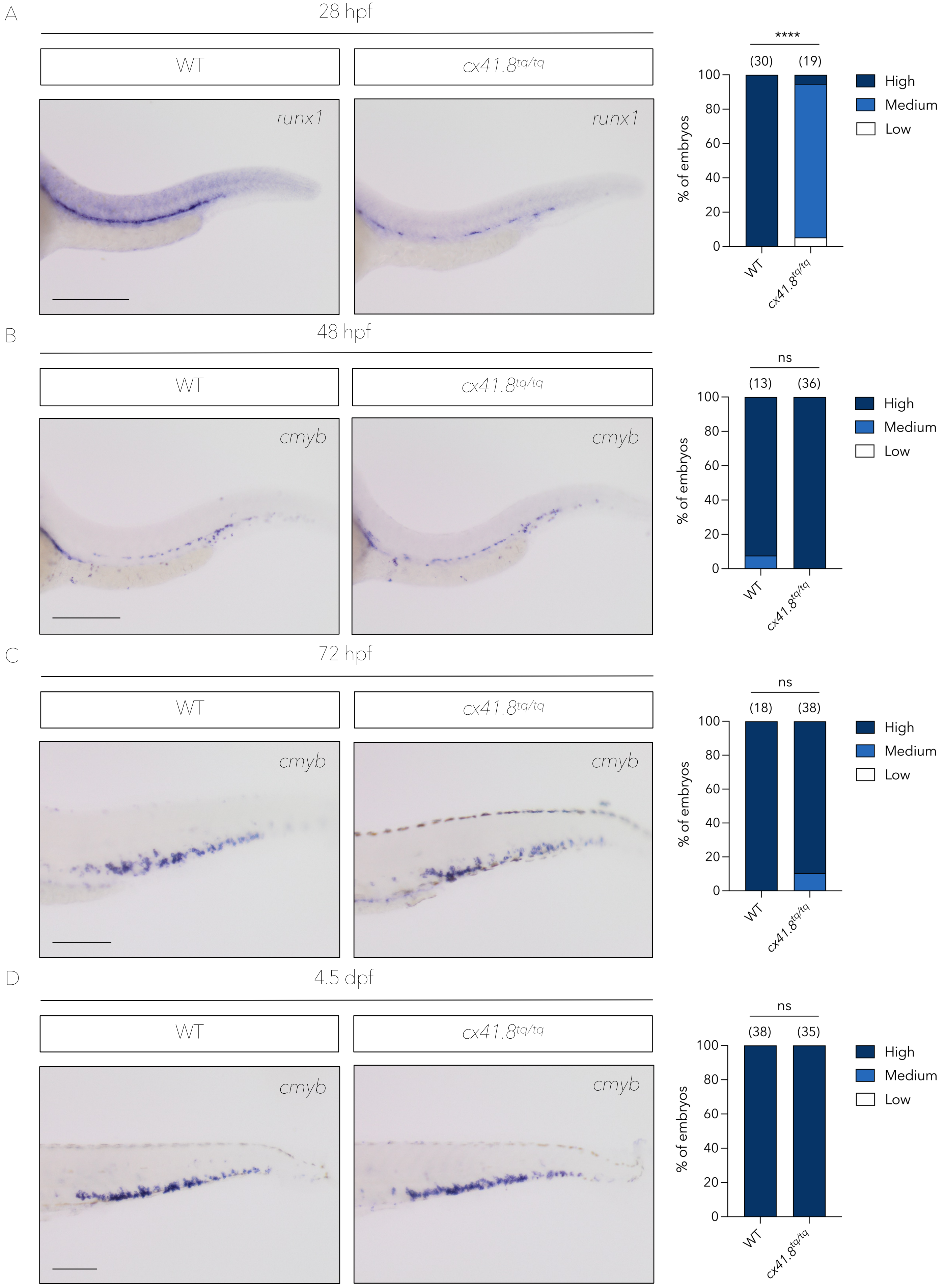

**Supplementary Figure 2. *cx41.8$^{tq/tq}$* mutant embryos display a delay in HSPC specification**

**A.** *runx1 in situ* hybridisation and quantification in *cx41.8$^{tq/tq}$* mutants and controls at 28hpf. **B.** *cmyb in situ* hybridisation and quantification in *cx41.8$^{tq/tq}$* mutants and controls at 48hpf. **C.** *cmyb in situ* hybridisation and quantification in *cx41.8$^{tq/tq}$* mutants and controls at 72hpf. **D.** *cmyb in situ* hybridisation and quantification in *cx41.8$^{tq/tq}$* mutants and controls at 4.5dpf. Statistical significance was calculated using either a Chi-squared test (**A**) or Fisher's test (**B**-**D**). *p < 0.05, **p < 0.01, ***p < 0.001, ****p < 0.0001. Scale bars: 200 µm (**A**, **B**, **C** and **D**).

Supplementary Figure 3

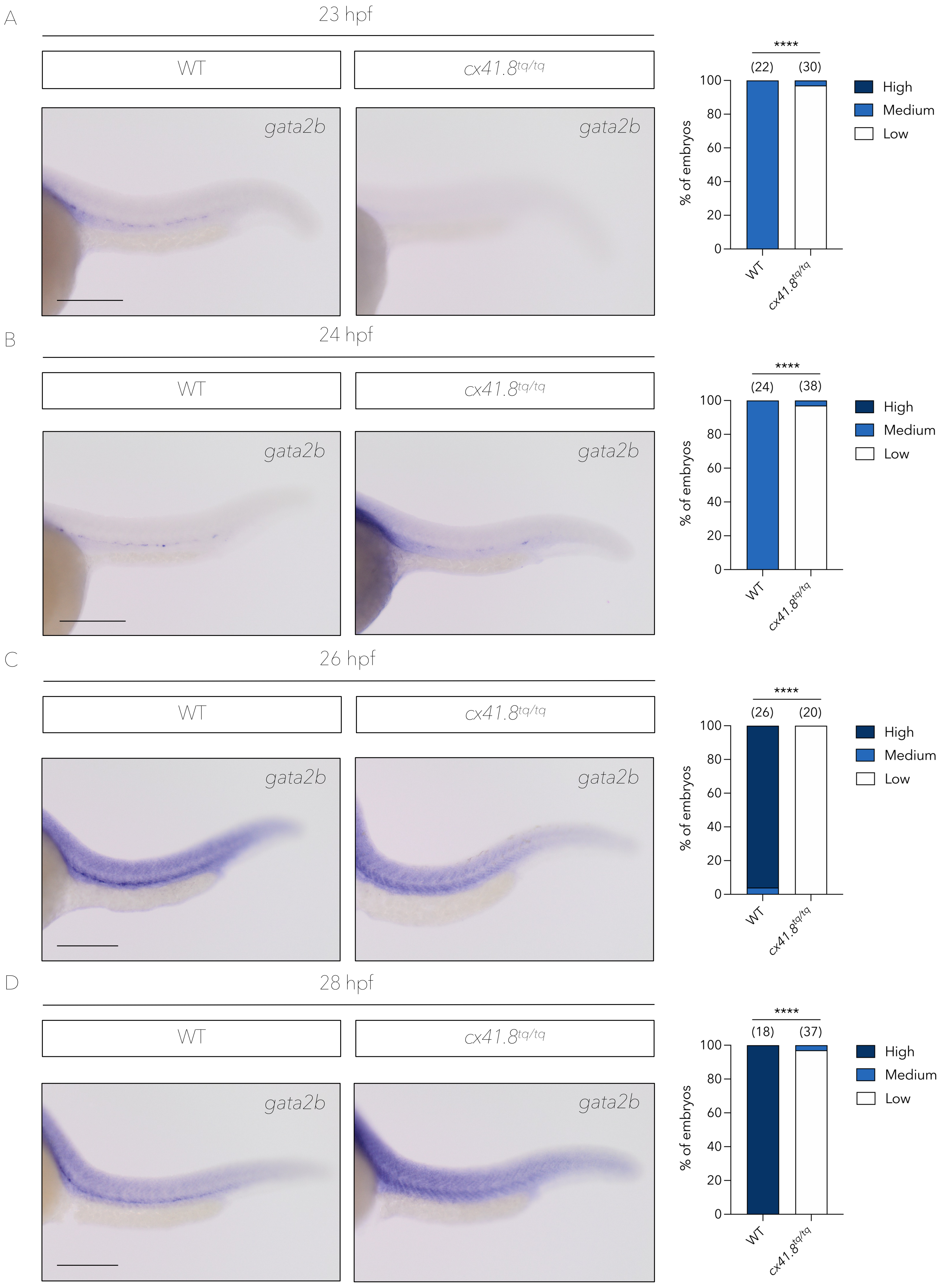

**Supplementary Figure 3. *cx41.8$^{tq/tq}$* mutant embryos display a delay in *gata2b* expression**

**A.** *gata2b in situ* hybridisation and quantification in *cx41.8$^{tq/tq}$* mutants and controls at 23hpf. **B.** *gata2b in situ* hybridisation and quantification in *cx41.8$^{tq/tq}$* mutants and controls at 24hpf. **C.** *gata2b in situ* hybridisation and quantification in *cx41.8$^{tq/tq}$* mutants and controls at 26hpf. **D.** *gata2b in situ* hybridisation and quantification in *cx41.8$^{tq/tq}$* mutants and controls at 28hpf. Statistical significance was calculated using either a Fisher's test (**A** and **B**) or Chi-squared test (**C** and **D**). *$p < 0.05$, **$p < 0.01$, ***$p < 0.001$, ****$p < 0.0001$. Scale bars: 200 µm (**A**, **B**, **C** and **D**).

Supplementary Figure 4

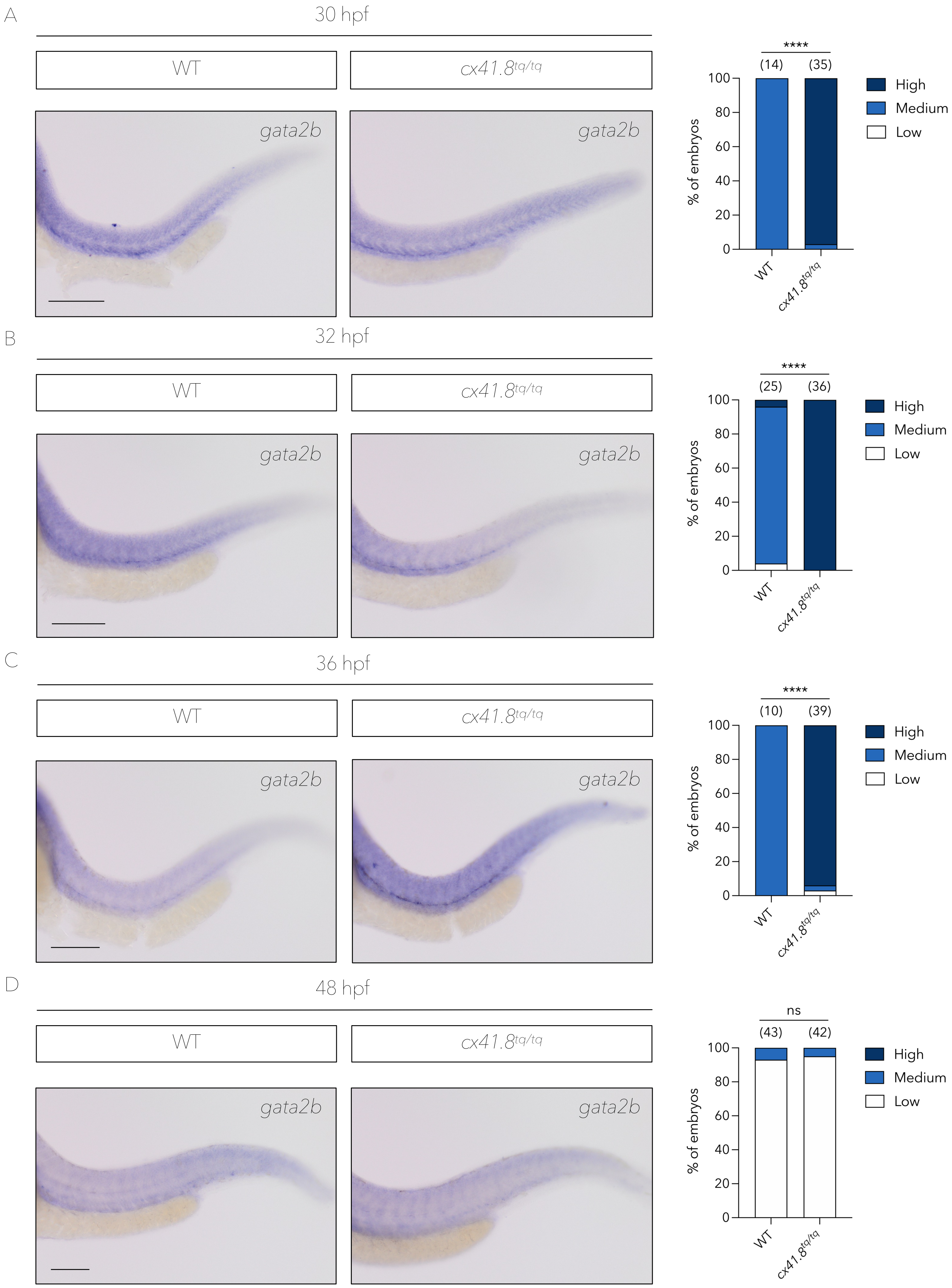

**Supplementary Figure 4. *gata2b* is expressed from 30hpf in *cx41.8*$^{tq/tq}$ mutant embryos**

**A.** *gata2b in situ* hybridisation and quantification in *cx41.8*$^{tq/tq}$ mutants and controls at 30hpf. **B.** *gata2b in situ* hybridisation and quantification in *cx41.8*$^{tq/tq}$ mutants and controls at 32hpf. **C.** *gata2b in situ* hybridisation and quantification in *cx41.8*$^{tq/tq}$ mutants and controls at 36hpf. **D.** *gata2b in situ* hybridisation and quantification in *cx41.8*$^{tq/tq}$ mutants and controls at 48hpf. Statistical significance was calculated using either a Fisher's test (**A** and **D**) or Chi-squared test (**B** and **C**). *p < 0.05, **p < 0.01, ***p < 0.001, ****p < 0.0001. Scale bars: 200 μm (**A**, **B**, **C** and **D**).

Supplementary Figure 5

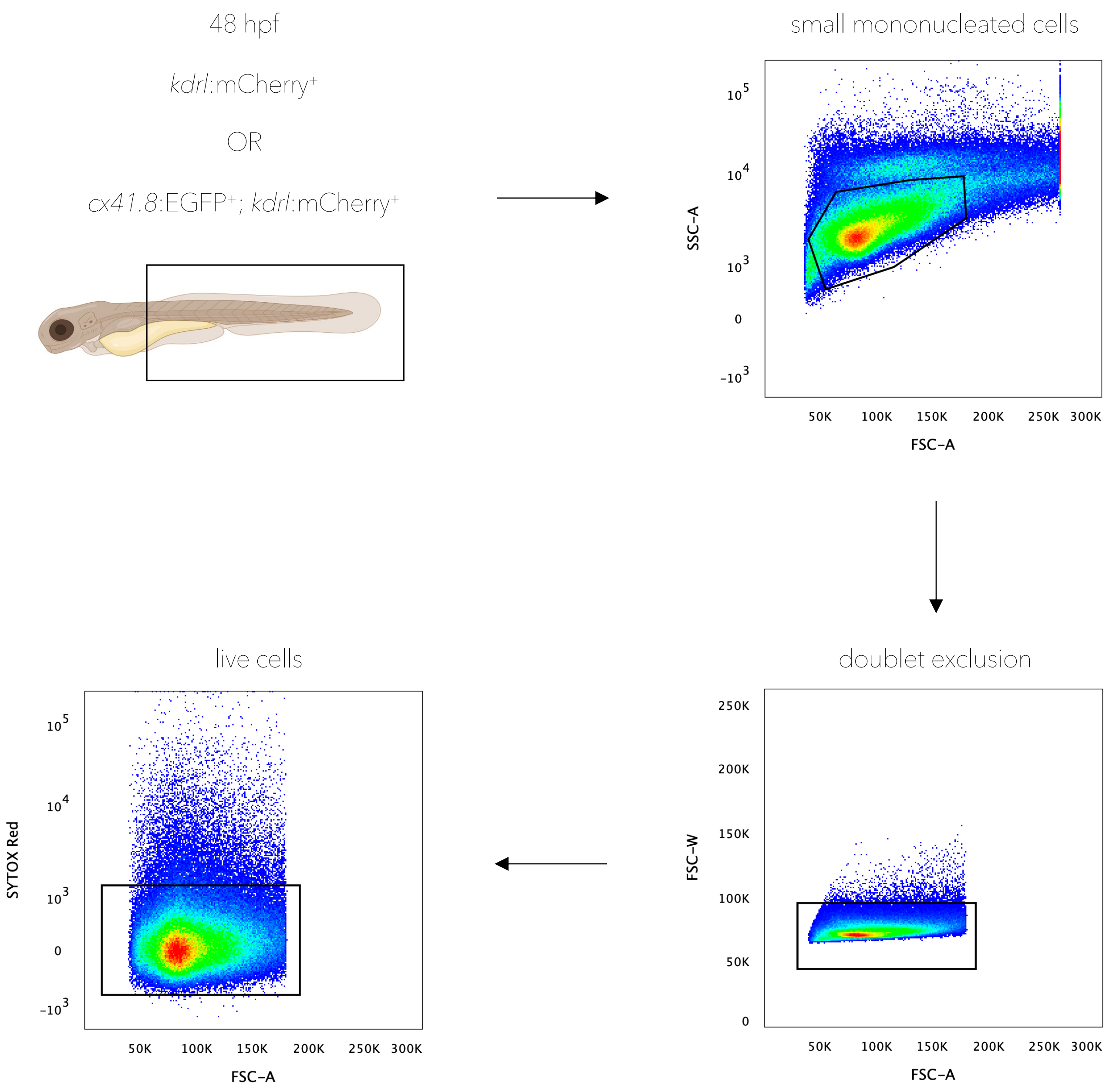

**Supplementary Figure 5. Gating strategy for flow cytometry analyses**

Trunk and tail dissection strategy and flow cytometry gating strategy for 48hpf *kdrl*:mCherry$^+$ or *cx41.8*:EGFP$^+$; *kdrl*:mCherry$^+$ embryos.

Supplementary Figure 6

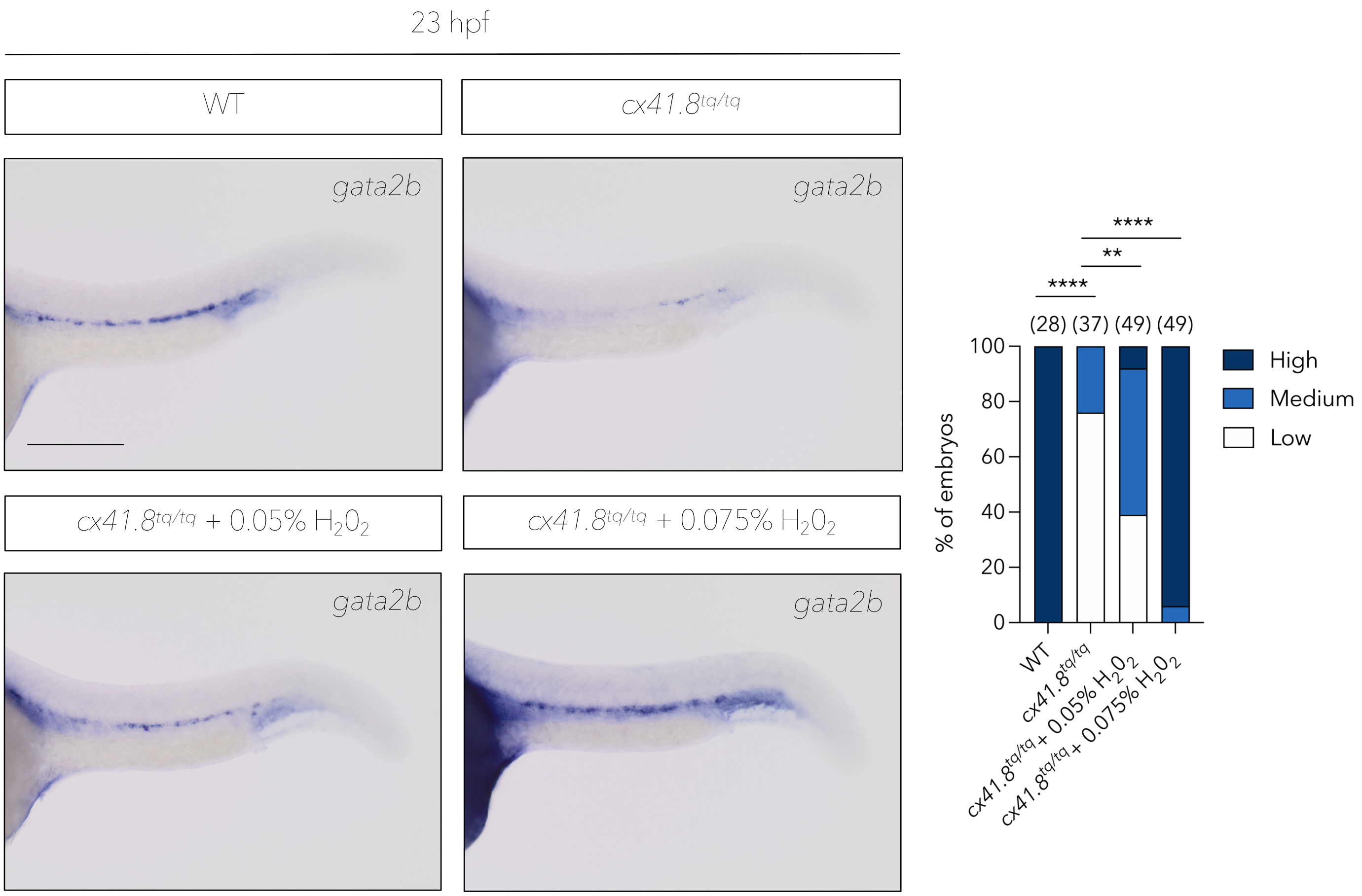

**Supplementary Figure 6. ROS induction in *cx41.8$^{tq/tq}$* mutant embryos rescues *gata2b* loss in a dose dependent manner at 23hpf**

*gata2b in situ* hybridisation and quantification in controls and *cx41.8$^{tq/tq}$* mutants supplemented with either 0.05% or 0.075% $H_20_2$. Statistical significance was calculated using a Chi-squared test. *$p < 0.05$, **$p < 0.01$, ***$p < 0.001$, ****$p < 0.0001$. Scale bar: 200 μm.

Supplementary Figure 7

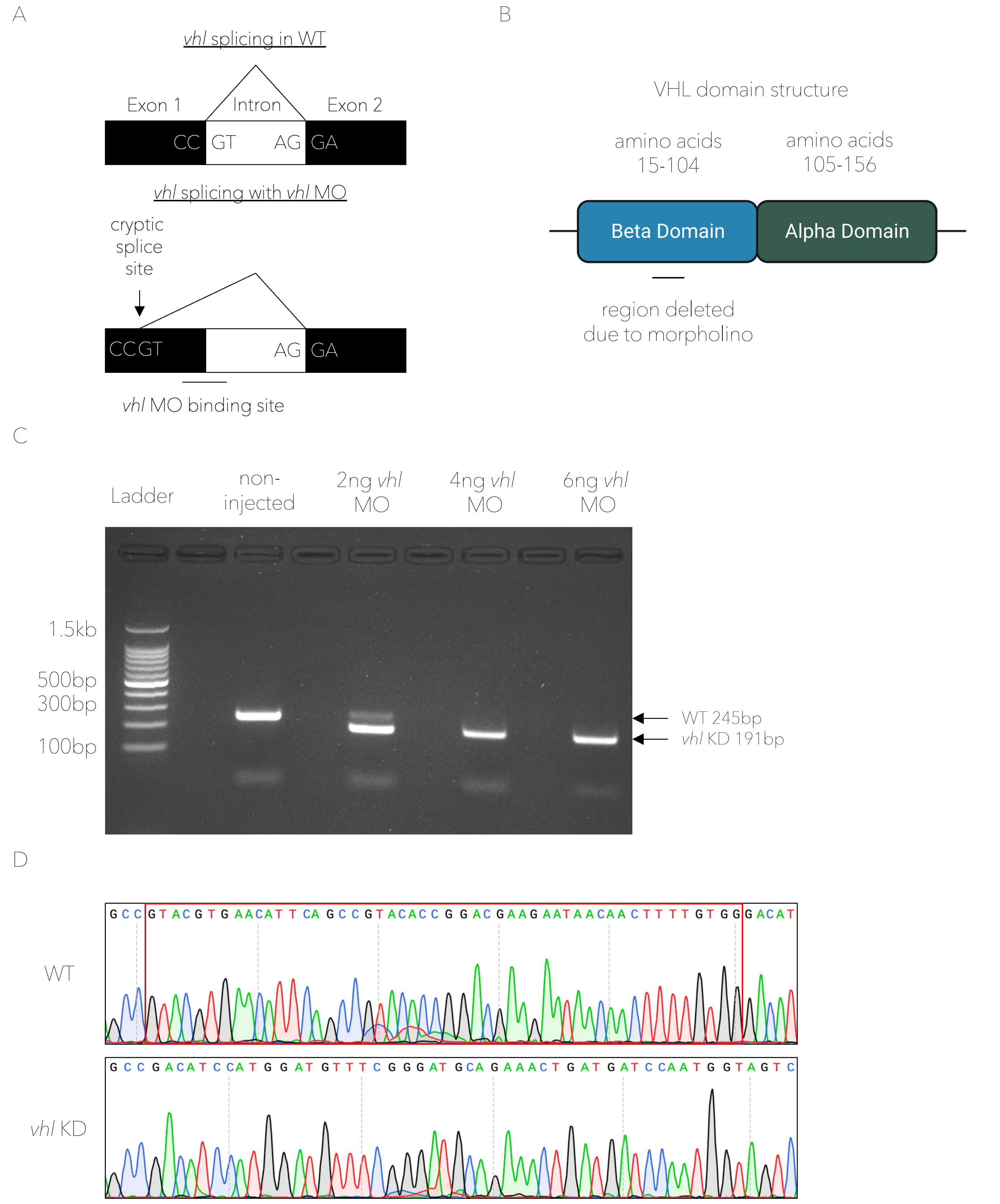

**Supplementary Figure 7. The *vhl*-MO results in the subsequent loss of 18 amino acids from the VHL beta domain**

**A.** Schematic to show the location of the *vhl*-MO induced cryptic splice site in exon 1 of the *vhl* transcript. **B.** Schematic to show the location of the *vhl*-MO induced loss of 18 amino acids from the VHL beta domain. **C.** Gel electrophoresis image showing the *vhl*-MO induced 54bp loss in the *vhl* transcript with 6ng of *vhl*-MO. **D.** Sanger sequencing confirming the 54bp loss in the *vhl* transcript with 6ng of *vhl*-MO. The red box in the WT sequencing track indicates the 54bp lost in the *vhl* transcript upon *vhl*-MO injection.

### Supplementary Figure 8

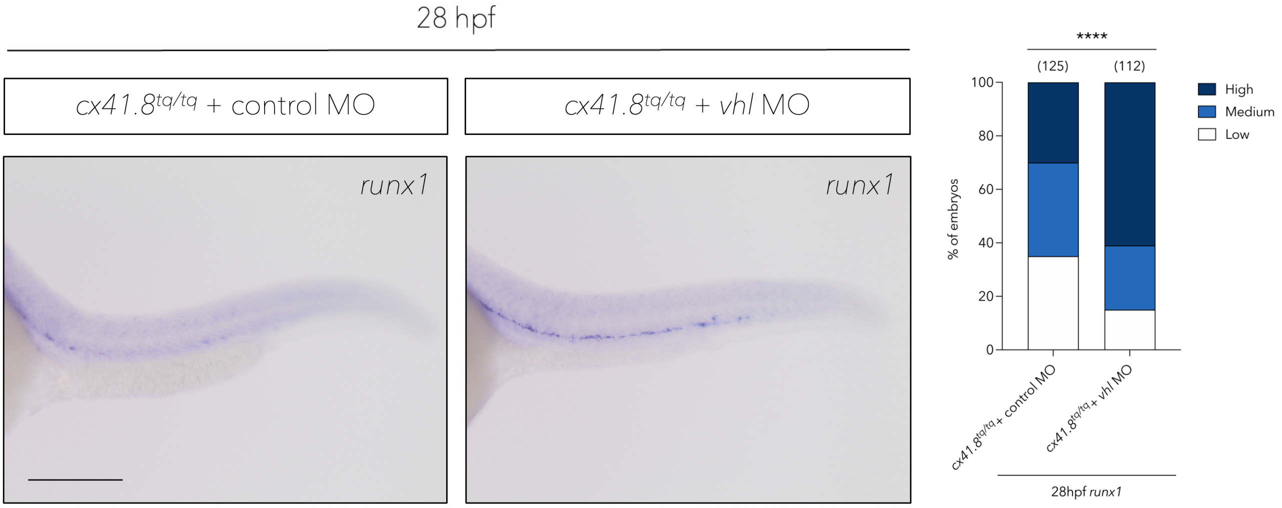

**Supplementary Figure 8. *vhl* knockdown rescues HSPC specification at 28hpf**

*runx1 in situ* hybridisation and quantification in control-MO or *vhl*-MO treated *cx41.8*$^{tq/tq}$ mutant embryos. Statistical significance was calculated using a Chi-squared test. *p < 0.05, **p < 0.01, ***p < 0.001, ****p < 0.0001. Scale bar: 200 µm.
